# Uncovering the Genetic Basis of Heterodichogamy in *Pterocarya* and *Cyclocarya* Using a Low-Input Pan-Genomic Approach

**DOI:** 10.1101/2025.03.10.642285

**Authors:** Hao-Sheng Liu, Wei-Hao Wang, Yan-Feng Song, Yang Yang, Da-Yong Zhang, Wei-Ning Bai, Bo-Wen Zhang

**Affiliations:** Ministry of Education Key Laboratory for Biodiversity Science and Ecological Engineering, College of Life Sciences, Beijing Normal University, 100875 Beijing, China; The National Facility Preservation Bank for Forestry and Grassland Germplasm Resources, Beijing Forestry University, Beijing, 100083, China

**Keywords:** Heterodichogamy, Mating system, Juglandaceae, *Pterocarya*, Structure Variation

## Abstract

The heterodichogamous mating system, characterized by two distinct mating types (protogyny and protandry), is rare among flowering plants, but it is present in nearly all species in Juglandaceae (the walnut family). Recent studies have identified distinct structural variations underlying heterodichogamy in *Juglans* and *Carya*. To verify the independent origins of this trait in Juglandaceae and investigate whether structural variations also drive heterodichogamy in *Juglans* closely related genera, we explored its genetic basis in *Pterocarya* and *Cyclocarya*. Using a pan-genome graph approach, we identified a structural variation region associated with mating types across the *Pterocarya* genus. This region includes 30 kb tandem repeats in the dominant allele and an insertion in the recessive allele, with shared polymorphisms spanning 78 kb from the 3′UTR of *S12e*, covering a *FAF-like* gene, to a *Gypsy* transposable element. Downstream analyses suggest that the specific expression of *FAF-like* gene and small RNAs uniquely expressed from the tandem repeats of dominant allele regulate heterodichogamy. Further investigation in *Cyclocarya* identified nine candidate loci associated with heterodichogamy, which are non-homologous regions with those found in *Pterocarya*, *Juglans*, and *Carya*. These findings provide novel evidence for the multiple independent originations of convergent genetic basis in regulating heterodichogamy in Juglandaceae and highlight the utility of pan-genome approaches in deciphering structural variation-associated traits.

## Introduction

Structural variations (SVs) are key drivers of phenotypic diversity in plants, governing a wide range of evolutionary processes from ecological adaptation to phenotypic differentiation^1–4^. With the recent advances in long-read sequencing, multiple studies have revealed the pivotal role of SVs in sex determination or regulating mating systems across diverse species. For instance, in *Pistacia vera*^5^ and in *Silene latifolia*^6^, SVs drive the evolution of sex chromosomes^5^, while in *Linum*, SVs were associated with distyly traits^7^. These findings collectively position SVs as fundamental drivers in the evolution of plant reproductive strategies.

Heterodichogamy, a mating system characterized by the temporal separation of male and female flowering phases (protogyny or protandry), promotes non-random mating by synchronizing reciprocal flowering times, reducing self-fertilization while enhancing outcrossing compared to typical dichogamous plants^8–10^ (Figure 1A). Though rare among flowering plants (documented in 13 families and 20 genera to date^11,12^), heterodichogamy exhibits remarkable conservation across Juglandaceae (the walnut family), where nearly all species display this trait^8,10,11,13,14^. Recent breakthroughs in *Juglans* and *Carya*, two genera of Juglandaceae, revealed two distinct ancient SVs controlling heterodichogamy with different associated genes^15^, suggesting either independent origins or ancestral origin with subsequent genetic turnover of this trait within the family. To resolve this evolutionary paradox and further determine whether SVs are the primary mechanism controlling heterodichogamy across Juglandaceae, further investigation into additional species is essential.

**Figure 1.**
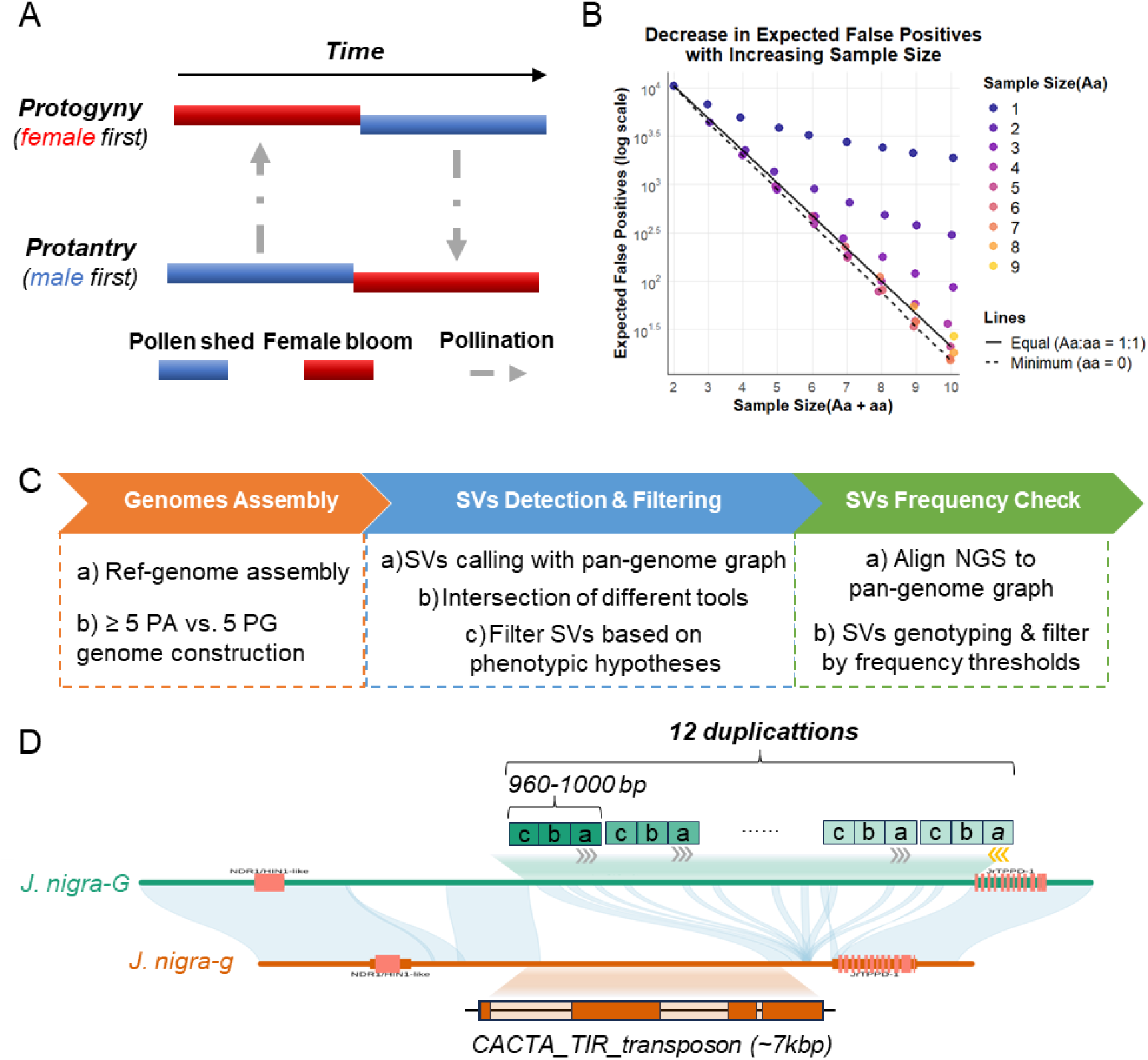
New pipeline detects heterodichogamy controlled by a single locus. (A) Diagram of floral phenology and pollination dynamics under heterodichogamy. Male-first (protandry) and female-first (protogyny) morphs exhibit synchronized but temporally reversed flowering phases. Gray arrows indicate pollen transfer directions between morphs. (B) Theoretical false-positive rate (*Pfp*) of structural variant detection under varying sampling strategies. (C) Three-step pipeline for SV prioritization in heterodichogamy studies. (D) Validation of the pipeline in *Juglans nigra* identifies four high-confidence SVs (Chr07:30.71–30.73 Mb), overlapping the known heterodichogamy locus.

In this study, we focused on *Pterocarya*, a genus within Juglandaceae closely related to *Juglans*^16^ and known to exhibit heterodichogamy (Figure S1). Unlike economically valuable *Juglans* and *Carya* species, which have extensive phenotypic data due to cultivation, *Pterocarya* and *Cyclocarya* species inhabit temperate forests^17–20^, making phenotypic data collection challenging. This limitation restricts the application of traditional genetic mapping approaches, such as GWAS (Genome-Wide Association Study), which typically require large numbers of phenotyped individuals. However, the genetic architecture of heterodichogamy offers a unique advantage for study. In *Juglans* and *Carya*, heterodichogamy is controlled by a single gene locus with two alleles (dominant and recessive), following Mendelian inheritance^14,21^. Additionally, structural variations have been shown to play a critical role in regulating this trait^15^. Leveraging these insights, we aimed to identify structural variations associated with heterodichogamy in *Pterocarya* and *Cyclocarya* by obtaining third-generation sequencing data from a small number of phenotyped individuals.

To achieve this, we developed a rapid pan-genomic pipeline that leverages the power of pan-genome analysis, which is particularly effective for detecting SVs associated traits in non-model organisms with limited sample sizes. By constructing a pan-genome graph based on a high-quality reference genome of *Pterocarya stenoptera* and resequencing a small number of individuals with known phenotypes, we aimed to pinpoint the structural variations associated with heterodichogamy. Through this approach, we identified a ∼78 kb structural variation that is distinct from the known control regions in *Juglans* and *Carya*. Furthermore, our transcriptomic analyses revealed that both *FAF* (*FANTASTIC FOUR*)*-like* gene and small RNAs derived from these SVs were involved in flowering regulation in *Pterocarya*. Additional implement of the pipeline in six *Cyclocarya* individuals also uncovered nine independent candidate heterodichogamy-loci in *Cyclocarya*, which are distinct from those in *Pterocarya*, *Juglans*, and *Carya*. These findings provide new insights into the genetic basis of heterodichogamy in Juglandaceae and demonstrate the efficacy of our pan-genomic approach in deciphering trait-associated structural variations in non-model organisms.

## Results

### A low-input genomic pipeline to decipher heterodichogamy

Unlike species of the *Juglans* and *Carya* genera, which have accumulated substantial phenotypic data, there are very few individuals of *Pterocarya* with accurately recorded flowering data. Additionally, Groh et al. found that the genetic loci associated with heterodichogamy in *Juglans* and *Carya* are related to longer segments of SVs^15^. Therefore, our research delineates a novel genomic analysis pipeline for studying species with limited phenotypic data.

To evaluate the feasibility of identifying heterodichogamy-associated SVs with limited samples, we modeled allele frequency distributions under the genetic architecture of heterodichogamy. Assuming single-locus Mendelian control and complete disassortative mating with two alleles^13^, we define protandrous (PA) individuals as homozygous recessive (*gg*) and protogynous (PG) individuals as heterozygous (*Gg*), reflecting the near absence of dominant homozygotes (*GG*) in natural populations. Under this model, a true causal locus controlling dichogamy morph would exhibit heterozygous genotypes (*Gg*) in all PG individuals and homozygous recessive genotypes (*gg*) in all PA individuals. However, a non-causal locus could spuriously match this pattern if, by chance, it also displays heterozygous genotypes (*Aa*) in all PG individuals and homozygous recessive genotypes (*aa*) in all PA individuals. This scenario would lead to false positives. For a biallelic neutral locus unrelated to the dichogamy trait, the probability (*Pfp*) of observing heterozygous genotypes (*Aa*) in *n* PG individuals and homozygous recessive genotypes (*aa*) in *m* PA individuals is given by:

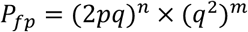

where *p* and *q* represent the frequencies of the dominant (*A*) and recessive (*a*) alleles, respectively, with p + q = 1. The terms 2*pq* and *q*^2^ represent the probabilities of observing the heterozygous genotype (*Aa*) and homozygous recessive genotype (*aa*), respectively. When *p* and *q* are directly derived from the observed genotypes in the sampled individuals (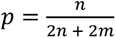 and 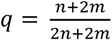), *Pfp* reaches its maximum value. This reflects the maximum possible false-positive rate under the given sampling conditions.

Under this scenario, we evaluated the expected genome-wide false-positive count (Figure 1B), which was scaled by an empirical genomic context observed in Juglandaceae (about 50,000 SVs according to our data in Juglandaceae, equivalent to one bi-allelic non-singleton SV >50 bp per 10 kb). When n = m = 5, *Pfp* reached 20.88, representing manageable false positives when combined with downstream filtering. Deviations from this 1:1 ratio (e.g., eight PA vs. two PG) elevated false positives by nearly 150-fold, while smaller total samples (n + m ≤ 8) generated prohibitive error rates (∼100 false SVs). A modest increase in PG samples (*n*) while holding total size constant further reduced *Pfp*, though the natural 1:1 PA/PG ratio in wild populations makes balanced sampling biologically and statistically optimal. This framework guided our experimental design: a minimum five PA vs. five PG sampling strategy balances field practicality with analytical rigor, as the resultant ∼20 false positives can be effectively eliminated through subsequent screening steps.

Guided by this theoretical framework, we designed a three-step pipeline to prioritize candidate SVs under Mendelian control and complete disassortative mating (Figure 1C). First, the pipeline begins with high-quality genome assembly of a minority of individuals (e.g., five PA vs. five PG) with known phenotype, followed by the enhancement of contig-level assemblies to pseudochromosome-level. Subsequently, a pan-genome graph is constructed by integrating these assemblies with a custom reference genome, enabling comprehensive capture of genomic diversity. Second, structural variations are detected and filtered, focusing on identifying variations that exhibit homozygosity in all recessive PA individuals and heterozygosity in most dominant PG individuals. Third, candidate structural variants underwent genotype frequency analysis using population-scale resequencing data, integrating both novel and public datasets devoid of phenotypic annotations. Genotype proportions were evaluated against the expected 1:1 ratio of heterozygous vs. homozygous, retaining variants exhibiting non-significant statistical divergence (χ² test, α=0.05) from the theoretical distribution while maintaining complete homozygosity in recessive PA individuals. Following the above three steps, we obtained the candidate structural variations associated with heterodichogamy.

To evaluate the performance of our pipeline, we applied it to *Juglans nigra* (black walnut), a species with a known heterodichogamous genetic region^15^. We first constructed a graph-based pan-genome through reference alignment (with five PGs and four PAs), identifying 70,098 structural variants (SVs >50 bp). At step two of the pipeline, we filtered candidates based on their genotypes: retaining only SVs that were heterozygous in PG individuals and homozygous in PA individuals. This initial filtering narrowed candidates to 35 SVs across 23 genomic regions. Subsequent population frequency analysis (n=62 resequenced individuals) further refined these to one genomic region with four high-confidence SVs (Chr07: 30.71-30.73 Mb), which were consistent to the previously characterized causal region (Figure 1D; Table S4). This application attests to the pipeline’s effectiveness and suggests its potential utility in investigating genetic variations in other species where phenotypic data is limited.

### Haplotype-resolved genome assemble of protogyny and protandry *Pterocarya stenoptera*

To better identify the genomic regions and candidate genes underlying the heterodichogamy in *Pterocarya*, we performed third generation long-reads sequencing and *de novo* assembly of *Pterocarya stenoptera*, which exhibit both protandry and protogyny (Pste-PA, Pste-PG). High-quality PacBio HiFi sequencing produced 63.66 Gb and 64.72 Gb of data per genome, yielding high coverage (111.69× and 113.55×; Table S1). *De novo* assembly generated contigs with high N50 values of 35.38 Mb (*Pste*-PA) and 35.94 Mb (*Pste*-PG) (Table S2). These assemblies were further scaffolded into chromosome-level genomes using high-throughput chromatin conformation capture (Hi-C) reads, providing phased haplotypes for comparative analysis.

The resulting consensus genomes (555.9 and 549.8Mb) showed size consistency with published Juglandaceae references while exhibiting superior continuity (≤ five gaps), representing a significant improvement compared to previously published genomes. The quality and completeness of the assemblies were evaluated using the Benchmarking Universal Single-Copy Orthologs (BUSCO)^15^ analysis, with values of 98.8% and 99.1% for the PA and PG genomes, respectively (Figure S1), and the long terminal repeat (LTR) assembly index (LAI)^16^ scores, which were also higher than 15 (Table S2), confirming the high quality of the genome assemblies.

### Identification of structural variations underlying heterodichogamy in *P. stenoptera*

In addition to the two individuals mentioned above for assembling the reference genomes, we sampled four additional PA and four PG individuals in an artificially planted population in Beijing, China and generated an average of 20.3 Gb of PacBio HiFi resequencing data (average depth: 35.5×). First, these data were *de novo* assembled into contigs, which were then scaffolded to pseudochromosomes by aligning them to the pre-constructed reference genome. Second, we constructed a graph-based pan-genome (∼847.08 Mb, containing 95,960 SVs) by integrating each genome from five protandrous and five protogynous individuals into the reference genome. To identify candidate SVs, we applied both minigraph^22^ and Panpop^23^ for SV detection, and retained only the intersection of their results to ensure high confidence. Then, we filtered candidates based on the genotypes: retaining only SVs that were heterozygous in all PG individuals and homozygous in all PA individuals, based on the observed rarity of homozygous PGs in *Juglans* (< 2% in the broad geographic sampling^15^). With this stringent filtering, we identified five candidate regions spanning Chr01, Chr03, Chr07, Chr09, and Chr06. However, despite the low probability, we further considered the possibility of one homozygous PG in our five samples. This expanded the candidate set to ten regions, including five additional loci on Chr03, Chr07, Chr08, Chr09, and Chr11. Third, to prioritize candidates, we tested whether SV genotypes in 96 short-read resequenced *P. stenoptera* individuals (unknown phenotypes) conformed to a 1:1 ratio. Of the ten candidate regions, seven exhibited highly significant deviations from the expected 1:1 ratio (χ² test, *P* < 0.003) and were excluded. Two of the remaining loci (Chr08: 29.086 Mb; Chr09: 25.299 Mb) showed marginal deviations (*P* < 0.03), leaving Chr06 (4.054-4.061 Mb) as the sole candidate with no statistical deviation (*P* > 0.068). Finally, we built up a regional tree using a total of 10 kb collinear regions nearby the identified SVs, which demonstrated a clear segregation of *G* and *g* haplotypes across the samples (Figure 2C).

**Figure 2.**
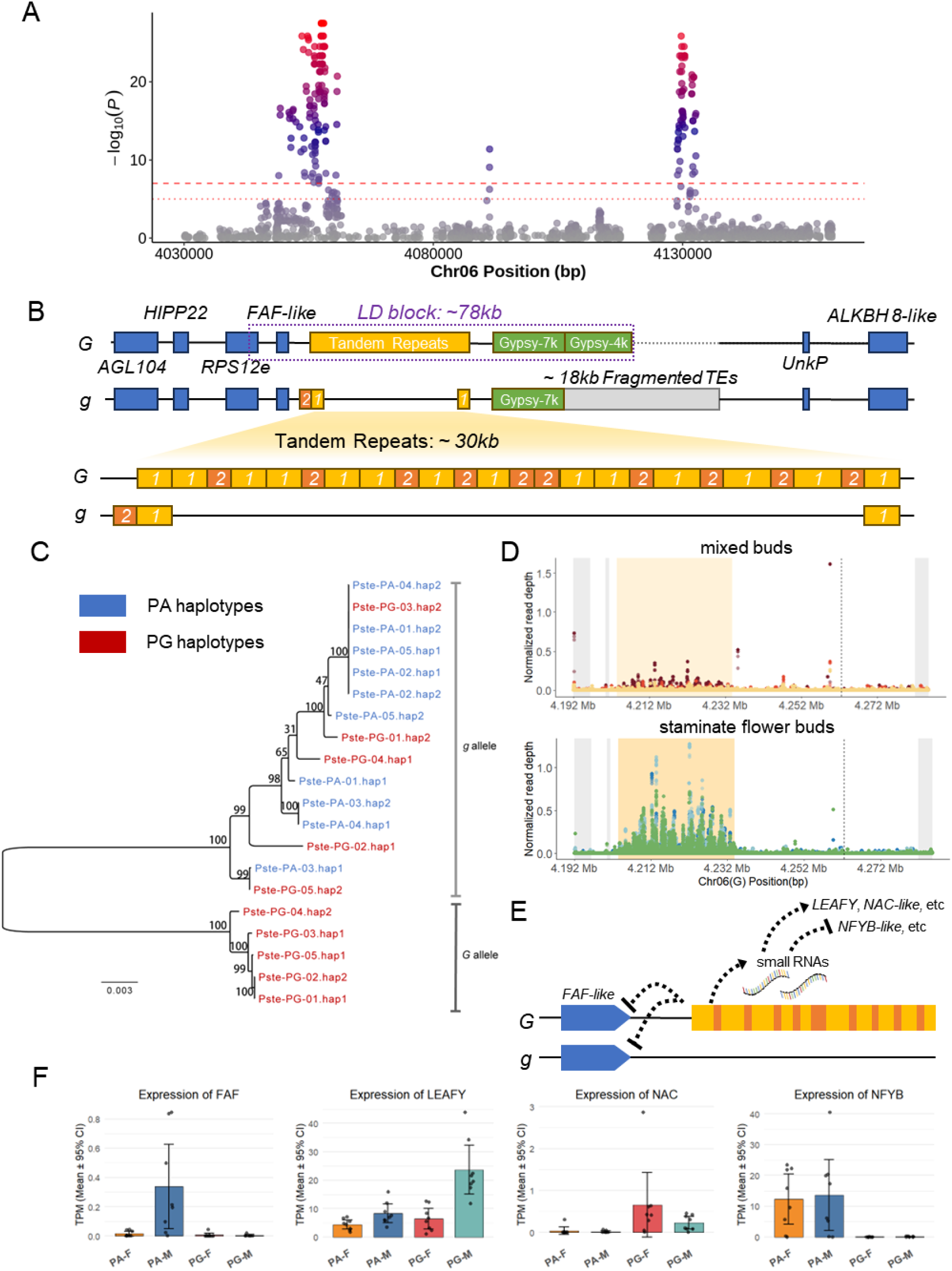
Structural and regulatory architecture of the heterodichogamy locus in *Pterocarya stenoptera.* (A) Locus Zoom Plots for the Genome-wide association study (GWAS) results using inferred protandrous and protogynous phenotypes. Dashed line marks summary statistics of *P* < 1×10^-5^ and 1×10^-7^. (B) Genomic structure of the *G* locus and structure of the *G*-allele-specific tandem repeats. Blue boxes denote flanking genes; yellow and orange boxes mark the tandem repeat region unique to the *G* allele: Unit-1 (1.6 kb, yellow); Unit-2 (600 bp, orange). Gene abbreviations: *AGL104* (agamous-like MADS-box protein 104), *HIPP22* (heavy metal-associated isoprenylated plant protein 22), *S12e* (ribosomal protein S12e gene), *FAF-like* (*FANTASTIC FOUR*-like), *UnkP* (an unknown function protein) and *ALKBH8-like* (alkylated DNA repair protein alkB homolog 8-like). (C) An unrooted maximum likelihood tree of 10 kb collinear regions from *G*/*g* haplotypes, with numbers on the branches indicating the bootstrap support values. Abbreviations: Pste (*P. stenoptera*), PA (protandry) and PG (protogyny). (D) Dot plots showing Tissue-specific small RNA expression from the *G* locus. *G*-allele repeats produce abundant sRNAs in staminate flower buds, while mixed buds show minimal expression. (E) Schematic plot summarizing Dual regulatory model: *G* allele suppresses the *FAF-like* homolog (*cis*-regulation, left) and generates sRNAs targeting *LEAFY-like*/*NAC-like* genes (*trans*-activation, right). Arrow: putative mobile sRNA signaling. (F) Bar plots showing the expression of *FAF-like*, *LEAFY-like*, *NAC-like*, and *NFYB-like* genes across tissues. *FAF-like* is PA-specific in staminate flower buds.

To conclusively validate the Chr06 candidate locus, we designed primers for Chr06 and two additional regions (Chr08 and Chr09) previously excluded by statistical filtering. PCR and Sanger sequencing of 18 phenotyped *P. stenoptera* samples (eight PAs and ten PGs) confirmed these findings: Chr06 SVs perfectly co-segregated with heterodichogamy (PA=*gg*, PG=*Gg*), while Chr08/Chr09 variants displayed discordant genotypes, conclusively rejecting their causality (Figure S2). The combined statistical rigor and experimental validation establish the SV around Chr06:4.054-4.061 Mb as the singular genomic locus associated with this developmental polymorphism.

### Characters of Structural Variations Associated with Heterodichogamy in *P. stenoptera*

To determine the size of the linkage region associated with the SV in *P. stenoptera*, we conducted a genome-wide association study (GWAS) using presumptive protandrous and protogynous phenotypes (Figure 2A). Phenotype assignment was based on validated SVs on Chr06, which served as genetic markers in the association analysis of 96 resequenced individuals. The associated SNPs delineated a single linkage region spanning 4.0536-4.1322 Mb (78 kb) on Chr06, hereafter referred to as the *G* locus (Figure 2B). The upstream boundary of the *G* locus lies within the intron of the ribosomal protein S12e gene (*S12e*), while the downstream boundary is located approximately 5.5 kb from the nearest gene, which encodes a protein of unknown function (*UnkP*). Notably, the locus harbors a *FAF-like* gene belonging to the *FANTASTIC FOUR (FAF)* gene family, which has been reported to regulate flower development in tomato^24,25^. We identified 15 *FAF* family members in *P. stenoptera*, including this *G*-locus *FAF-like* gene. The conserved domain (PF11250) is present across all members, supporting its classification within the *FAF* family (Figure S5A). Comparative analysis of the coding region between *G* and *g* haplotypes revealed 43 segregating SNPs (25 of which were non-synonymous), indicating deep allelic divergence (Figure S5B).

Within the linkage region of *G* haplotype, a ∼30 kb tandem repeat is located approximately 2 kb downstream of the *FAF-like* gene (Figure 2B). This repeat region consists of two main units: Unit-1 (∼1,000 to 2,100 bp, averaging ∼1,600 bp), which contains a palindromic Mutator TIR transposon (∼380 bp), and Unit-2 (∼400 to 670 bp, averaging ∼600 bp), organized in an alternating tandem array of 14 Unit-1 and eight Unit-2 sequences (Figure 2B). Approximately 16.7 kb downstream of this repeat region, the *G* haplotype contains two distinct *Gypsy* transposable elements (TEs, 7 kb and 4 kb, respectively), marking the end of the linkage region. In contrast, the *g* haplotype harbors only a single copy of (Unit-2 + Unit-1) and a single Unit-1, located 2 kb downstream of the *FAF* gene, with a ∼29 kb sequence annotated as fragmented TEs between the two copies. Additionally, the *g* haplotype retains the 7 kb *Gypsy* found in the *G* haplotype, followed by a ∼18 kb region containing other fragmented TEs.

To explore potential gene-structure associations, we annotated genes proximal to the linkage regions (Figure 2B). Starting approximately 17 kb upstream of the linkage region, we identified the agamous-like MADS-box protein 104 (*AGL104*) and heavy metal-associated isoprenylated plant protein 22 (*HIPP22*) genes. The left boundary of the linkage region encompasses the terminal three coding sequences (CDS) and the 3’ untranslated region (3’ UTR) of the *S12e* gene, although no SNPs associated with two alleles were detected within its coding regions. Downstream of the linkage region, at distances of approximately 5.5 kb and 19 kb, we identified genes encoding a protein of unknown function (*UnkP*) and an alkylated DNA repair protein alkB homolog 8-like (*ALKBH8-like*), respectively.

### Transcriptomic Insights into the Mechanism of Structural Variation-Mediated Heterodichogamy

To explore the function of the *G* locus, we performed mRNA and small RNA sequencing on staminate flower buds and mixed buds. Comparative transcriptomic analysis between PA (*gg* carriers) and PG (*Gg* carriers) individuals identified 469 differentially expressed genes (DEGs) in staminate flower buds and 325 DEGs in mixed buds (Figure S6, Table S5). Among these DEGs, the *FAF-like* gene within the *G* locus exhibited significantly increased expression (logFC = -6.22, FDR < 0.0001) in PA staminate flower buds (Figure 2F), while no significant differences in mixed buds. As for the other five genes (*AGL104*, *HIPP22*, *S12e*, *UnkP* and *ALKBH8-like*) near the *G* locus, no significant differences were found in either staminate flower buds or mixed buds (Figure S3). Notably, the *FAF-like* gene was exclusively expressed in PA staminate flower buds, with no detectable expression in PG staminate flower buds (Figure 2F) or other tissues (Figure S7). This pronounced expression pattern was not observed for the 14 other homologous *FAF* genes in *P. stenoptera* genome (Table S5), highlighting the unique role of the *FAF-like* gene within the *G* locus.

We identified small RNAs transcribed from the *G*-allele-specific repetitive regions (Figure 2D and S4). To assess their potential regulatory functions, we aligned these small RNAs across the genome and identified 1,209 putative target genes. These matches were distributed across coding sequences (191 genes), intronic regions (681 genes), and gene-proximal regions (362 genes within ±500 bp of gene boundaries). Integrating these findings with the DEGs revealed that 18 of the matched genes differentially expressed in staminate flower buds, six of which were also differentially expressed in mixed buds. Among the 18 staminate flower bud DEGs, nine genes showed increased expression in PG, while the remaining nine genes showed suppressed expression (Figure 2F and S8; Table S5). Notably, three of these genes (*NFYB-like*, *LEAFY-like* and *NAC-like*) have been implicated in the regulation of flower development^26–28^. Their differentially expression observed in PG (*Gg* carriers) indicate the potential *trans*-regulation role of the small RNAs from *G* allele-specific repetitive regions (Figure 2E).

### The Evolutionary Origin of the Identified Locus in the Genus *Pterocarya*

To determine whether the *G* locus in *P. stenoptera* is conserved across other species within the genus, we analyzed 119 resequenced individuals from four additional species (*P. macroptera*: 40, *P. rhoifolia*: 20, *P. fraxinifolia*: 30 and *P. hupehensis*: 29). Firstly, we examined whether the SNPs identified at *G* locus in *P. stenoptera* (89 SNPs with GWAS p < 1 × 10^-18^) were also present in these species. Among the 89 SNPs, six SNPs were shared across all four species. Notably, the genotype ratios of heterozygotes to recessive homozygotes based on these six SNPs closely matched the expected 1:1 equilibrium ratio (Figure 3A). Furthermore, the proportion of dominant homozygotes (PGs) across 119 *Pterocarya* individuals was consistently low (average 2.79%), matching the ratios reported in *Juglans*^15^.

**Figure 3.**
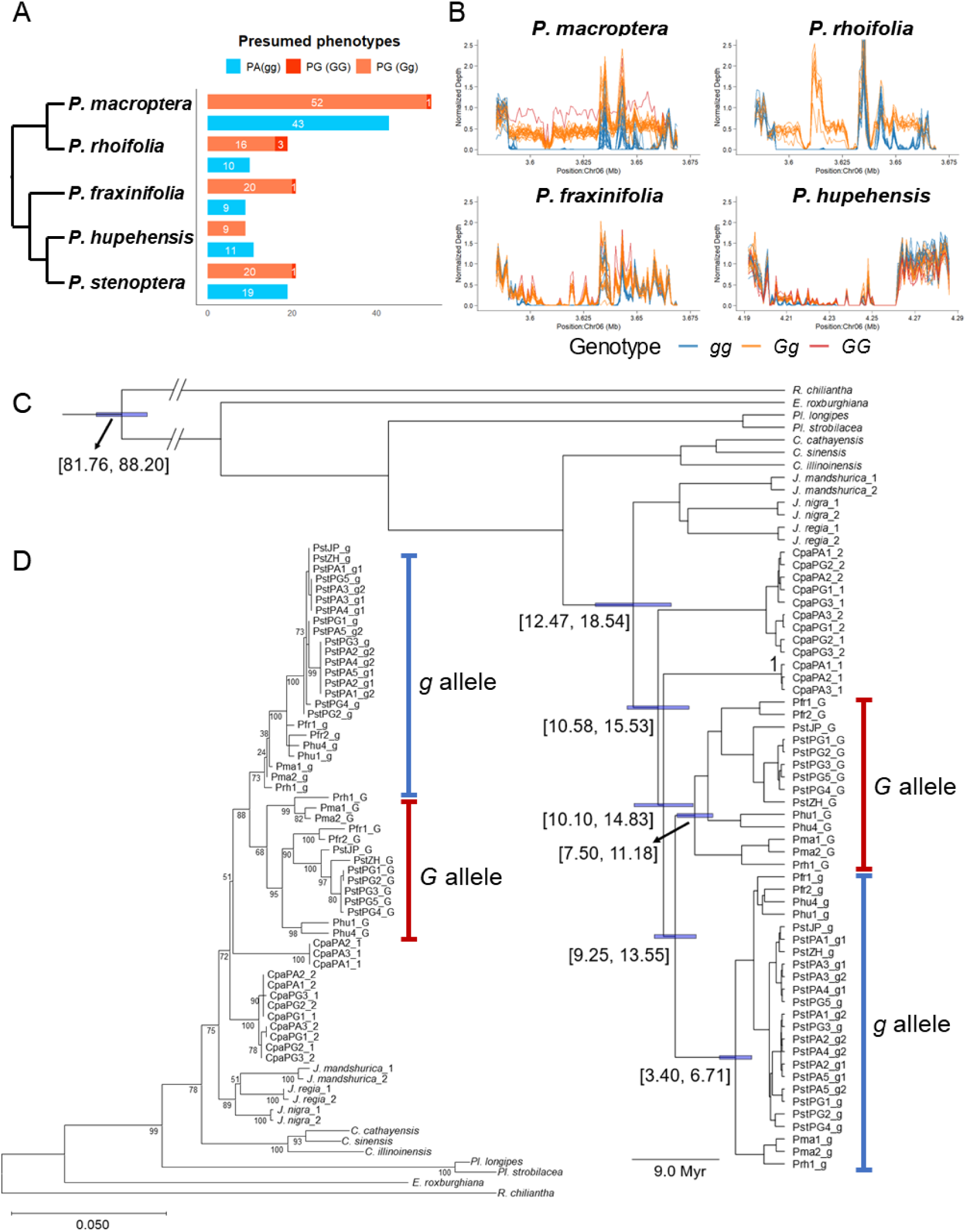
Evolutionary conservation and divergence of the heterodichogamy locus across *Pterocarya*. (A) Phylogenetic relationships and genotype ratios of five *Pterocarya* species. Left: Species tree (branch lengths scaled to divergence time). Right: Observed genotype frequencies align with a 1:1 equilibrium. Abbreviations: PA (protandry) and PG (protogyny). (B) Normalized read depth across the *G* locus in 119 individuals from four *Pterocarya species*. Alignment to *P. macroptera* references confirms three distinct genotypes (*GG*, *Gg* and *gg*). (C) Bayesian tree of 1,398 bp collinear regions from 12 Juglandaceae species and *Pterocarya G* and *g* haplotypes. Blue bars mark the 95% HPD Divergence date for the main clades. Species abbreviations: Pma (*P. macroptera*), Pst (*P. stenoptera*), Pfr (*P. fraxinifolia*), Prh (*P. rhoifolia*), Phu (*P. hupehensis*), Cpa (*Cyclocarya paliurus*). (D) Maximum-likelihood tree of the same 1,398 bp collinear regions from 12 Juglandaceae species. Species abbreviations: Pma (*P. macroptera*), Pst (*P. stenoptera*), Pfr (*P. fraxinifolia*), Prh (*P. rhoifolia*), Phu (*P. hupehensis*) and Cpa (*Cyclocarya paliurus*).

To further validate the structural conservation of the *G* locus across *Pterocarya* species, we mapped resequencing data from the 119 individuals to reference genomes of PG *P. stenoptera*. If the *G* locus governs mating types across the genus, inferred PA individuals (classified by diagnostic SNPs) should exhibit near-zero read depth in the tandem duplication region of the *G* allele, whereas inferred PG individuals (heterozygous *Gg* or homozygous *GG*) should show proportional depth increases. Remarkably, alignment patterns from the reference genomes revealed clear separation of the three genotypes (Figure 3B), consistent with the genotypic classifications derived from the shared SNPs. These findings provide robust evidence for the structural conservation of the *G* locus across the genus, supporting its role as a key determinant of heterodichogamy in *Pterocarya*.

To infer the evolutionary history of the *G* locus within *Pterocarya*, we constructed a regional tree using a total of 1,398 bp SV-free collinear regions within the *G* locus across 12 Juglandaceae species (Figure 3C and 3D). The maximum likelihood (ML) tree revealed a clear divergence between *G* and *g* haplotypes across all *Pterocarya* species. Bayesian molecular dating further estimated the divergence time of the *G* and *g* haplotypes to be approximately 9.25-13.55 million years ago (Myr) (Figure 3C and 3D), predating the estimated origin of *Pterocarya* (∼7.00 Myr)^29^. The phylogeny also indicated that the divergence of these haplotypes occurred more recently than the divergence of *Pterocarya* from its closely related genera, *Cyclocarya* and *Juglans*, suggesting an independent origin of the *G* locus in *Pterocarya*.

### Pan-genomic pipeline identifies different candidate loci for heterodichogamy in *Cyclocarya paliurus*

To complete the understanding of the heterodichogamy locus within Juglandeae, we expanded our pipeline to *Cyclocarya paliurus*, a genus sister to *Juglans* and *Pterocarya*. For this species, we generated haplotype-resolved assemblies of protandrous (Cpa-PA) and protogynous (Cpa-PG) morphs. We constructed a pan-genome for *C. paliurus* by integrating three PA vs three PG individuals, resulting in a total pan-genome size of ∼756.39Mb and identifying 90,139 SVs). Using our established SV screening pipeline, we identified 568 phenotype-associated SVs (Table S6). This number is consistent with the prediction of our statistical model (Figure 1B), suggesting that the limited sample size (n + m = 6) precluded the unambiguous identification of candidate loci. Filtering through resequencing of 40 individuals (35 unknown phenotype individuals and five known mating-type) controls refined the candidate list to nine SVs (Figure 4A). Among these, three SVs exhibited high probability due to increased numbers of segregated SNPs nearby the identified SVs (Figure 4B, 4C, S9 and S10). Notably, all nine candidate loci in *C. paliurus* were located in genomic regions distinct from the heterodichogamy control loci previously identified in *Pterocarya*, *Juglans*, and *Carya* (Figure 4A). This emphasize that heterodichogamy evolved independently in these four genera, rather than through inheritance of a shared ancestral system.

**Figure 4.**
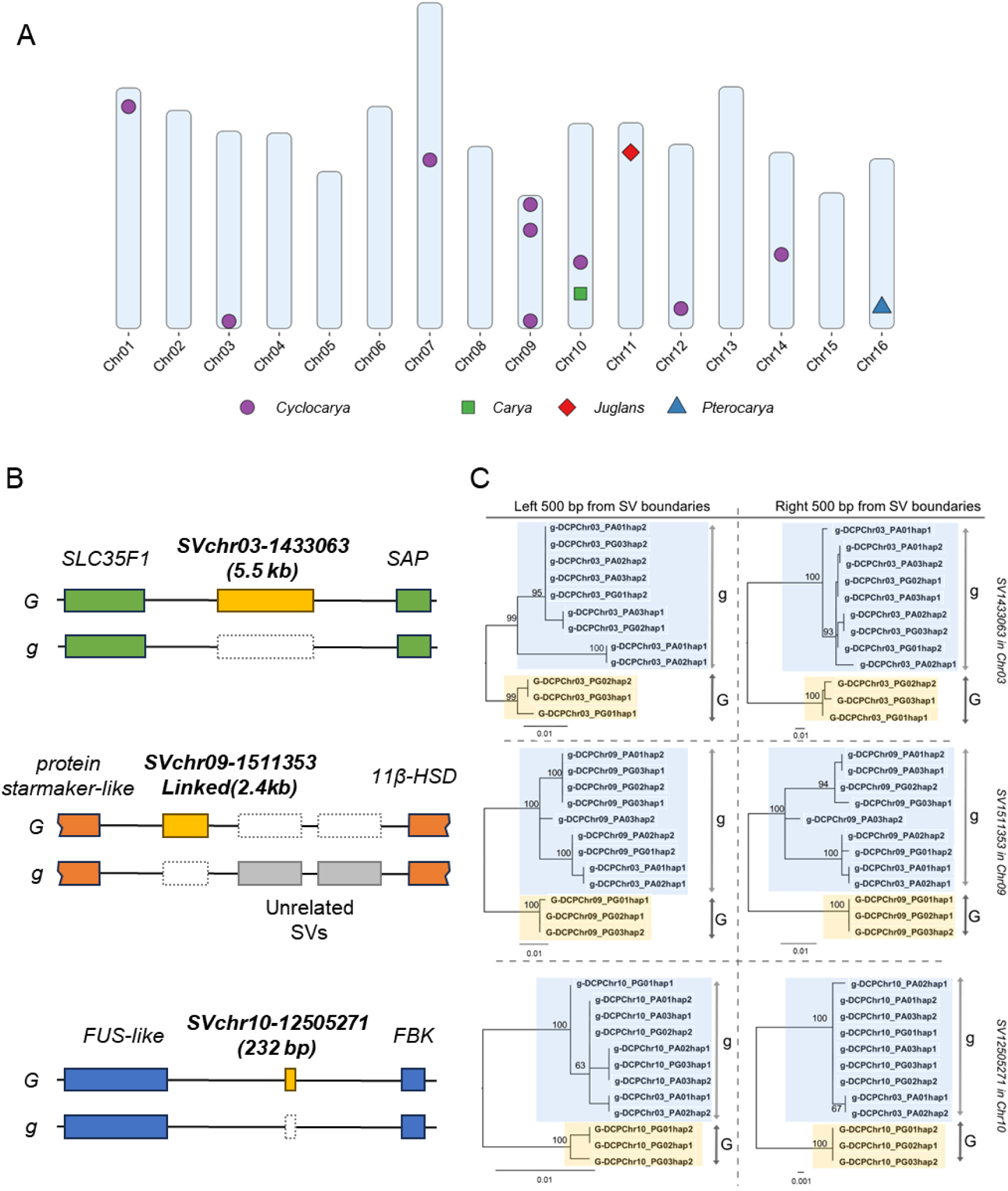
Genomic Divergence of Heterodichogamy-Associated SVs in *Cyclocarya paliurus* and Related Genera. (A) Genomic distribution of heterodichogamy-associated SVs across four genera. Candidate loci in *Cyclocarya paliurus* (purple circles) are spatially distinct from those in *Carya* (green squares), *Juglans* (red diamonds), and *Pterocarya* (blue triangles). (B) Structural organization of three candidate SV in *C. paliurus*. Gene abbreviations: *SLC35F1*(Solute carrier family 35 member F1-like), *SAP* (zinc finger A20 and AN1 domain-containing stress-associated protein), *11β-HSD* (11-beta-hydroxysteroid dehydrogenase), *FUS-like* (RNA-binding protein FUS-like) and *FBK* (F-box/kelch repeat protein). (C) Haplotype-resolved phylogeny using 500 bp sequence flanking the candidate SVs. Abbreviations: DCP (diploid *C. paliurus*).

## Discussion

Having successfully located the heterodichogamy loci in *Pterocarya*, as well as revealed candidate loci in *Cyclocarya*, our low input pan-genome pipeline has demonstrated several advantages over traditional approaches. First, it requires only a small number of phenotypic individuals, making it ideal for species like *Pterocarya* where large-scale phenotypic data collection is unfeasible. Second, it efficiently refined candidate variants by using allelic frequency obtained from phenotypic-unknown resequencing data. Third, pan-genome-based SV genotyping speeds up analysis and improves the resolution for the identified structural variations, which in the meantime, avoided the individual variances on the SV boundaries confounding the loci detection. Notably, such inheritance pattern is not unique for heterodichogamy, several studies on sex determining loci^6,30^ and distyly-associated loci^7,31^ has illustrated similar characteristics as heterodichogamy loci found across Juglandaceae. This makes our pipeline highly effective for studying genomic features where structural variations have been strongly linked with steadily phenotypic segregation.

The *G* locus coordinates protogynous flowering through dual *cis*- and *trans*-regulatory mechanisms. In PG individuals, the presence of *G* allele completely suppressed the expression of the adjacent *FAF* homolog in staminate flower buds, with no allele-specific expression from *g* allele in *Gg* carriers (Figure 2F). Given *FAF*’s conserved role in promoting floral meristem determinacy^24,25^, its downregulation likely delays male phase onset. Concurrently, *G*-specific small RNAs from the SV region target developmental regulators, including *LEAFY-like* genes that show elevated expression in PG male buds (Figure 2F). While mixed bud transcriptomes lack *G*-locus-proximal DEGs, we hypothesize that these mobile sRNAs may prime early female maturation through *LEAFY* activation—a scenario supported by *LEAFY*’s established function in accelerating floral fate commitment^27^. This dual regulatory architecture enables the *G* allele to coordinate both delayed male phase (via *FAF* suppression) and advanced female phase (via sRNA-mediated *LEAFY* activation), satisfying the developmental prerequisite for protogyny. Such antagonistic control of opposing floral phases through a single locus parallels evolutionary trajectories observed in other reproductive systems where trade-offs enhance outcrossing efficiency^7,32^. Future temporal transcriptomics across flowering stages could test whether sRNA gradients mediate developmental phase-shifting—a potential universal mechanism for heterodichogamy maintenance.

Although the controlling loci of heterodichogamy in four genera from Juglandaceae are non-homologous, the genetic architectures in *Juglans* and *Pterocarya* exhibit shared hallmarks: both *G* alleles contain tandem repeats composed of TE-derived sequences (a few hundred to a thousand bp units), suggesting TE-mediated repeat expansion. In *Juglans*, these repeats are tightly linked to the 3’ UTR of *TPPD1*. Similarly, in *Pterocarya*, a structurally analogous repeat array, potentially formed by TE insertion, is positioned approximately 2 kb downstream of the *FAF-like* gene and harbors multiple TE families. This pattern supports a model where SVs establish conserved genomic frameworks by suppressing recombination^33^, while TEs drive lineage-specific innovation through epigenetic and structural modifications^34–36^. In Juglandaceae, this functional synergy achieves both evolutionary stability and regulatory flexibility. SVs maintain haplotype integrity by preventing recombination, ensuring the persistence of heterodichogamy. Concurrently, TE-derived repeats within these SVs provide regulatory raw material, enabling fine-tuning of flowering time and floral development. In future studies, it will be interesting to dissect how TE-derived repeats influence gene regulation and how SVs maintain haplotype stability, which may help unravel the genomic mechanisms underlying the origins and maintenance of sexual diversity.

Taken together, our findings in combination with earlier studies uncovered the independent genetic loci controlling heterodichogamy in *Pterocarya* and *Cyclocarya* in Juglandaceae. The convergent emergence of TE-mediated insertion and repeat expansion constantly found in heterodichogamy species, as well as in the distyly species of *Linum*, *Primula* and *Turnera*, suggested the fundamental roles of SVs in the origin and evolution of the diverse reproductive strategies in plants. Our study not only motivate further work to illustrate the underneath molecular mechanisms that may underlie the convergent origin and maintenance of those SVs, but also provided a useful pipeline to uncover similar cases in other species.

## Materials and Methods

### Plant materials and genome sequencing

To dissect the genetic architecture of heterodichogamy in Juglandaceae, we collected individuals with confirmed dichogamy morphs (protandry and protogyny) for long-read sequencing and additional resequencing data from broader populations (phenotypes uncharacterized) across three species: *Pterocarya stenoptera*, *Cyclocarya paliurus* and *Juglans nigra*.

For *P. stenoptera*, 18 individuals from the National Olympic Forest Park (Beijing, China) were selected for pan-genome construction. Field observations during spring 2023 and 2024 classified their dichogamy morphs. PacBio HiFi sequencing (Sequel II) generated 11-64 Gb reads per individual (19.5-113× coverage), complemented by Hi-C and transcriptome data (Illumina NovaSeq 6000, PE150) for scaffolding. Population resequencing included 215 individuals spanning five *Pterocarya* species: *P. stenoptera* (96 individuals), *P. macroptera* (40), *P. rhoifolia* (20), *P. fraxinifolia* (30), and *P. hupehensis* (29), sequenced to 30× depth on Illumina NovaSeq 6000 (Table S3).

In *C. paliurus*, a pan-genome was assembled from six individuals in Hubei Province, China, using PacBio HiFi reads (35-143× coverage) and Hi-C scaffolding (Illumina NovaSeq 6000, PE150). Genetic diversity analysis leveraged resequencing data from 40 wild individuals across southern China (Illumina NovaSeq 6000 or DNBSEQ-T7, 30× depth).

For *J. nigra*, we selected 10 individuals in Beijing (two) and Gansu (eight), China, combining PacBio HiFi sequencing (37-128 coverage) with Hi-C data. Population genomic analyses integrated 62 individuals: 53 publicly available datasets (NCBI SRA: SRP149991; CNGB: CNP0001209)^37,38^ and nine newly sequenced samples from Gansu (DNBSEQ-T7, 30× depth).

### Genome assembly and quality assessment

The consensus sequences of *P. stenoptera*, *C. paliurus* and *J. nigra* were initially assembled. Hifiasm (v0.19.9-r616)^39,40^ was employed to generate primary contigs using PacBio HiFi and Hi-C reads, with each species utilizing protogyny Hi-C data. Subsequently, gfatools were utilized to convert these results into linear genomes, presented in fasta format. Hi-C data were further employed for scaffolding purposes using the Juicer^41^ and 3D-DNA^42^ tools. The scaffolding outcomes were manually optimized with JuiceBox^41^, which involved the rectification of chromosomal boundaries, resolution of misjoins, and handling of inversions and translocations, culminating in the derivation of 16 corrected pseudo-chromosomes. The original reads were employed to reduce genomic gaps via the quarTeT^43^ GapFiller approach. Chromosomal orientations were ascertained through comparative analysis with existing genomes.

To assemble haplotype-resolved genomes, the ‘Hi-C integration’ mode in Hifiasm^39,40^ was used to construct contig-level assemblies for both haplotypes by leveraging Hi-C and PacBio HiFi reads. Subsequently, phased contigs were aligned to the previously constructed consensus genome using RagTag^44^ for scaffolding purposes.

Genome quality was assessed using BUSCO^15^ (v5.5.0) with the embryophyta_odb10 dataset to evaluate completeness. Long terminal repeat (LTR) assembly index (LAI) scores were calculated using a custom script based on outputs from EDTA v2.0.1^45^, providing a measure of genome continuity and repeat region accuracy.

### Construction of Graph-based pan-genome and SV identification and SVs calling

To address the sequence representation limitations caused by collapse events when using linear reference genomes, we employed haplotype-resolved genomes for the construction of a graph-based pan-genome. Minigraph^22^ (v0.20-r559) was used to build the graph-based pan-genome, while Panpop^23^ was utilized for SV calling from small-sample PacBio long-read datasets, using default parameters.

For population-level SV genotyping, the variation graph tool vg^46^ (v1.58.0) was employed with default settings. Short reads from each accession were mapped to the graph-based pan-genome using the *vg giraffe* module, and SV genotyping results in VCF format were generated using *vg call*.

### Validations of Heterodichogamy-related SNPs and SVs

To validate the presence of candidate SNPs and SVs in populations and their association with the trait, samples of *Pterocarya* with known phenotypes were analyzed. Primers were developed for three initial candidate SVs, accounting for the high likelihood of PG samples being heterozygous. Validation was performed by examining whether the target fragments were successfully amplified and whether the amplification sequences corresponded to the observed phenotypes. This process identified a single candidate SV in *Pterocarya* as associated with the trait.

### Prediction and annotation of genes and repeats for the identified loci

Gene structures within the identified loci were extracted from the structural annotation files of the published *Pterocarya stenoptera* genome^47^. To functionally characterize the genes located in the vicinity of the loci, we performed sequence homology searches using the NCBI BLAST^48^ and structural predictions via the AlphaFold Protein Structure Database^49^. Transposable elements were annotated *de novo* using EDTA v2.0.1^45^, a comprehensive pipeline for TE identification and classification.

### Comparison of genetic structures between *G*/*g* alleles

We compared *G* (protogynous) and *g* (protandrous) allele architectures using phased assemblies processed through samtools (v1.19)^50^. Alignments were performed in MUMmer4^51^ with nucmer using k-mer=20 and mismatch penalty=-5 to detect structural variations. To minimize false alignments, delta-filter retained only one-to-one mappings (-m) with ≥90% identity (-i 90) and ≥ 100 bp length (-l 100). Mummerplot generated PDF-formatted synteny maps from filtered alignments.

### Genome-wide association analysis and Trait-associated loci hitch-hiking test

Genome-wide association studies (GWAS) were performed using PLINK^52^ v1.90. For *Pterocarya stenoptera*, the “DOM” model (dominant genetic effect model) in the --model function was selected based on the basic assumptions of heterodichogamy. Fisher’s exact test p-values were used to identify significantly associated variants.

To address potential hitchhiking effects at loci associated with heterodichogamy identified in smaller samples and further determine the range of the *G* locus, a “presumptive-phenotype GWAS” was conducted. Presumed phenotypes were determined based on genotypes linked to the trait and subsequently incorporated into the GWAS workflow. Manhattan and Q-Q plots were generated using the R package CMplot^53^.

### Phylogenetic reconstruction and the estimation of divergent times for the identified locus

Based on the results of collinearity analysis for SNP linkage region identification, we extracted the highly conserved regions within the 78-kb linkage block detected for the *G*/*g* alleles, manually cutting out all the SV and nearby regions and concatenated the remains. Then we used MAFFT with the iterative refinement method (the G-INS-I option) to rebuild the alignments (see Supplemental data 1). This process resulted in a total length of about 10,009 bp orthologous alignment for 10 vs 10 *Pterocarya* haplotypes, which includes the regions from the 3’ UTR of *S12e* to the front of Tandem Repeats and a small collinear segment after the Tandem Repeats up to the front of *Gypsy*. To obtain the alignment across 63 sequences from 12 Juglandaceae species, gaps and largely non-collinear sequences induced by additional species were deleted, which resulted to a collinear segment of a total length of 1,398 bp.

We then inferred the gene trees using Maximum likelihood method under the JC+FO model with 100 bootstraps using iqtree v2.3.6^54^. To implement the Bayesian molecular dating, we constructed Bayesian tree for the alignments from 12 Juglandaceae species using beast v2.7.7^55^ with HKY+estimated frequencies substitution model and Yule tree model to validate the gene tree and estimate the divergent date of the two haplotypes. We referred the oldest fossil of *Rhoiptelea*^56^ and set a prior distribution for the divergence of Juglandaceae following a log-normal distribution (M=4.43, S=0.02, offset=1.0) with a mean of 85 Myr and a 95% credible interval ranging from 81.7 to 88.3 Myr. The MCMC was run for 10^7^ iterations and burn-in for 10^6^ iterations. Samples were recorded every 1,000 iterations. We then used tracer to check and ensure all estimated parameters resulted conservation with ESS > 1,000.

### Transcriptomes analysis

For mRNA sequencing, total RNA was used to construct libraries. Poly(A) RNA was enriched, fragmented, and reverse-transcribed into cDNA, followed by adapter ligation and PCR amplification. Libraries were sequenced on the Illumina platform, generating paired-end reads with a minimum depth of 20 million reads per sample. Raw reads were preprocessed using fastp v0.23.4^57^ for adapter trimming and quality control. In order to better analyze the functional differences between *G* and *g* allele, newly assembled PG genomes were used as reference for comparative analysis. Clean reads were aligned to the reference genome using HISAT2^58^ with default parameters for accurate mapping. Read counts for each gene were generated using featureCounts^59^. Differential expression analysis was performed using edgeR v4.2.0^60^, with data normalization via the Trimmed Mean of M-values (TMM) method. The likelihood ratio test was applied to identify differentially expressed genes, with a significance threshold of |log2FC| > 1 and false discovery rate (FDR) < 0.05. For small RNA sequencing, total RNA was used to prepare libraries. Small RNAs were ligated with 3’ and 5’ adapters, followed by cDNA synthesis and PCR amplification. Libraries were size-selected (18-40 bp) and sequenced on the Illumina platform, generating at least 10 million reads per sample. raw reads were preprocessed using fastp v0.23.4 for adapter trimming and quality control. The processed reads were then independently mapped to both haplotype-level assemblies (male-first and female-first reference genomes) with the following parameters: zero mismatches were permitted, while allowing up to 12 distinct mapping locations per read to accommodate the repetitive structure of our target insertion regions.

Alignment files were processed using samtools v1.19^50^ to generate position-specific depth profiles. To enable cross-sample comparisons, we normalized the depth data using a depth-per-million scaling approach. For each sample, we first calculated the total sequencing depth by summing depth values across all genomic positions. A normalization factor was then derived by dividing 1,000,000 by the total depth. This factor was applied to each position’s depth value, generating normalized measurements that maintain the original genomic coordinates. The resulting normalized depth values represent depth units per million total depth units at each genomic position, facilitating quantitative comparisons across samples.

To investigate potential target genes regulated by small RNAs specifically produced in the staminate flower buds *G* allele region, small RNA reads consistently identified across all three replicates were extracted as query sequences. These queries were then subjected to BLAST searches against whole genome and CDS sequence databases using a word size of seven nucleotides. Potential target genes were identified by evaluating significant BLAST matches within CDS sequences and in both the gene body and its upstream and downstream regions (±500 bp), suggesting possible regulatory interactions.

## Acknowledgments

We thank Yi-Gang Song, Rui-Min Yu, Wei-Ping Zhang, Yu Cao, Ya-Mei Ding, Shi-Fu Xue, and Wan-Hua Lu for sample collection; Jian-Quan Liu, Yi Zhang, Ying Zhang, and Er-Li Pang for scientific discussions and helpful comments; This work could not have been completed without their support.

## Author Contributions

W.N.B., B.W.Z., and D.Y.Z. conceived and designed the project; H.S.L., W.H.W., Y.Y., Y.F.S., W.N.B., and B.W.Z. collected materials; H.S.L. and W.H.W. performed the bioinformatics analysis and PCR validation; W.N.B., B.W.Z., H.S.L. and W.H.W. wrote the paper. All authors approved the final version.

## Supplementary Material

**Figure S1.**
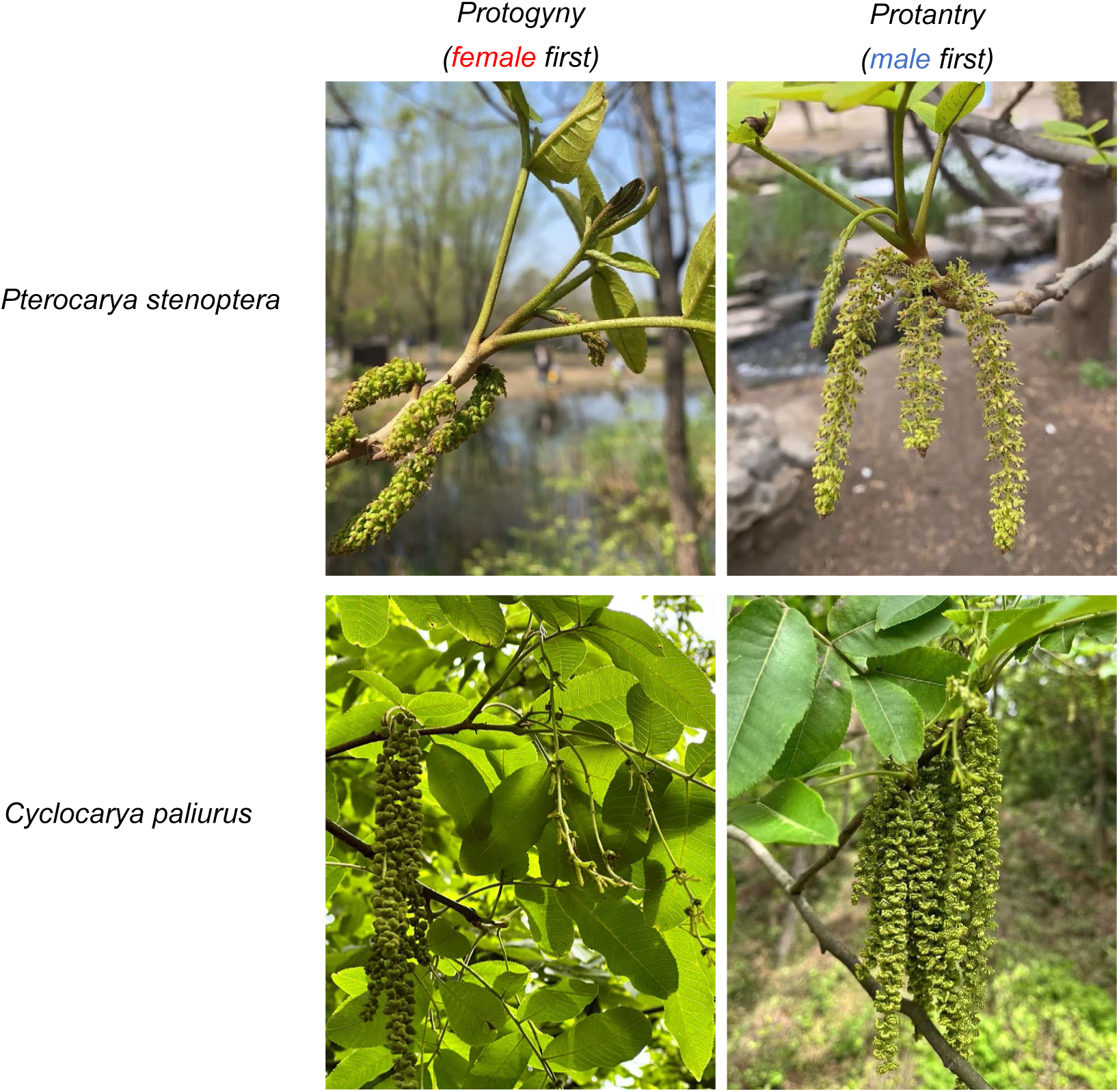
Female-first and Male-first individuals of *Pterocarya stenoptera* and *Cyclocarya paliurus*. Photo by H.S.L. and Y.F.S.

**Figure S2.**
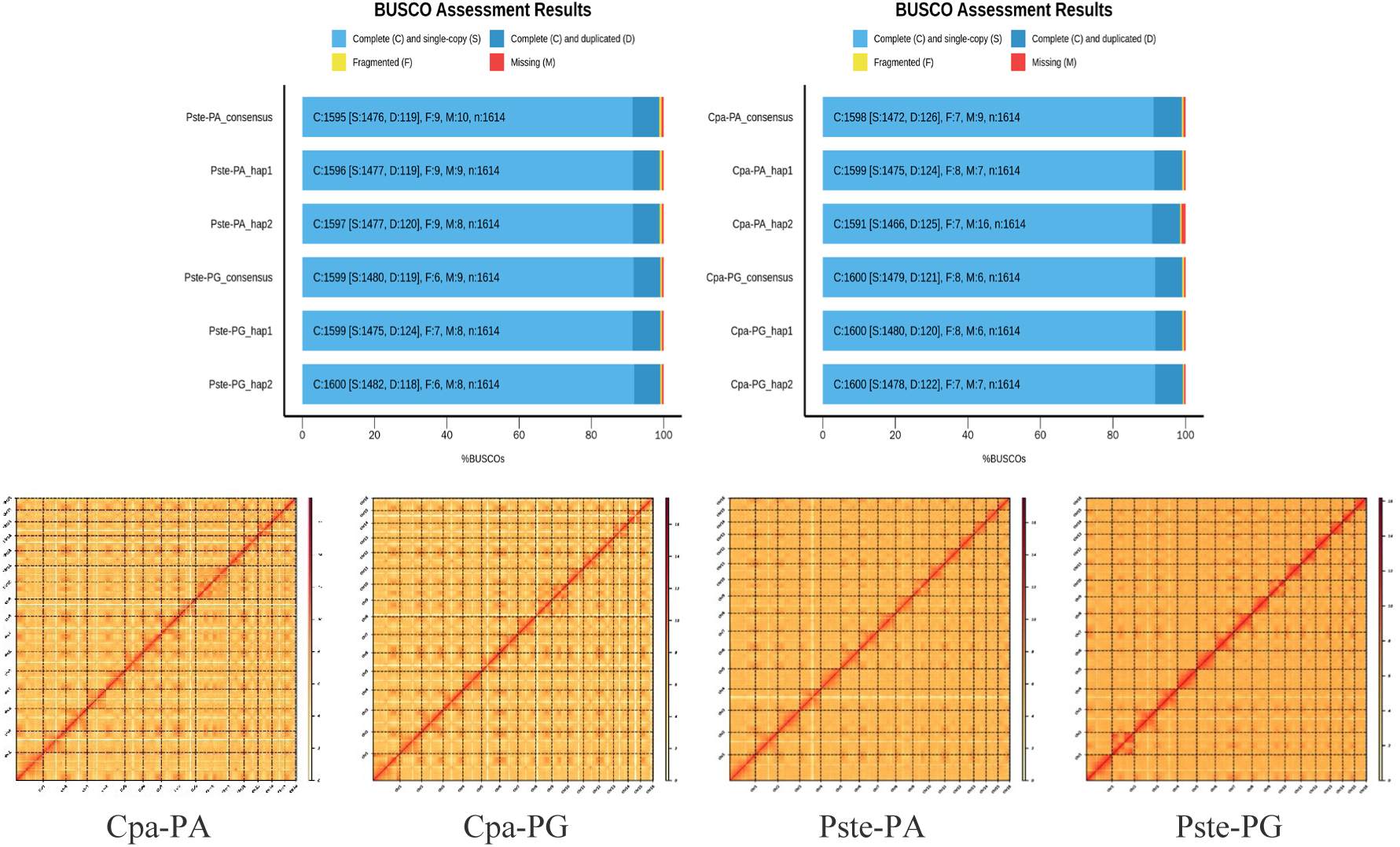
Summary of consensus and haplotype-resolved genome assembly. **Hi-C heatmap and BUSCO plot for four consensus genomes and corresponding hap1 & hap2 genomes.**

**Figure S3.**
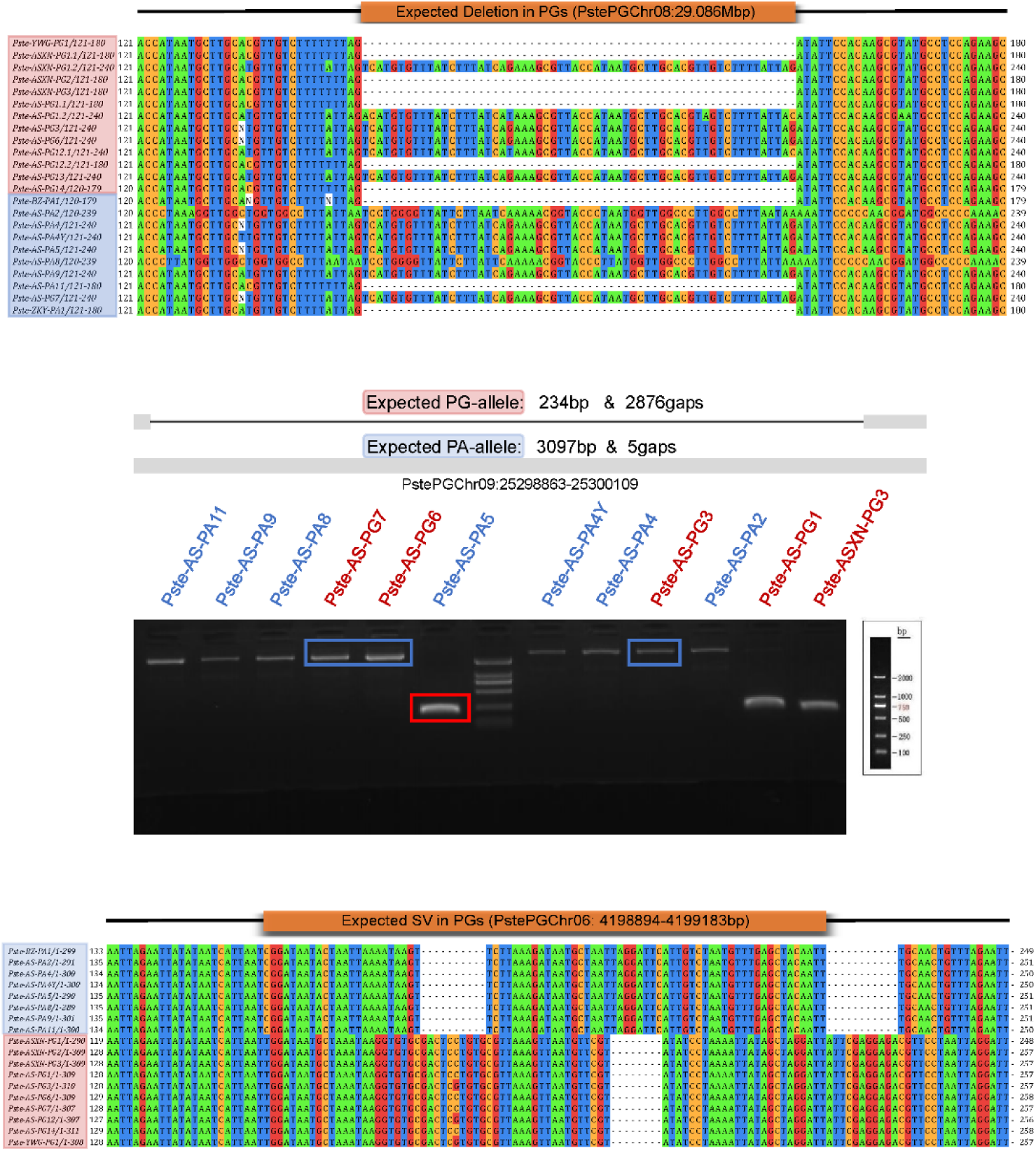
PCR validation of candidate SVs. PCR validation negated candidates on Chr08 and Chr09, further corroborating the candidate on Chr06.

**Figure S4.**
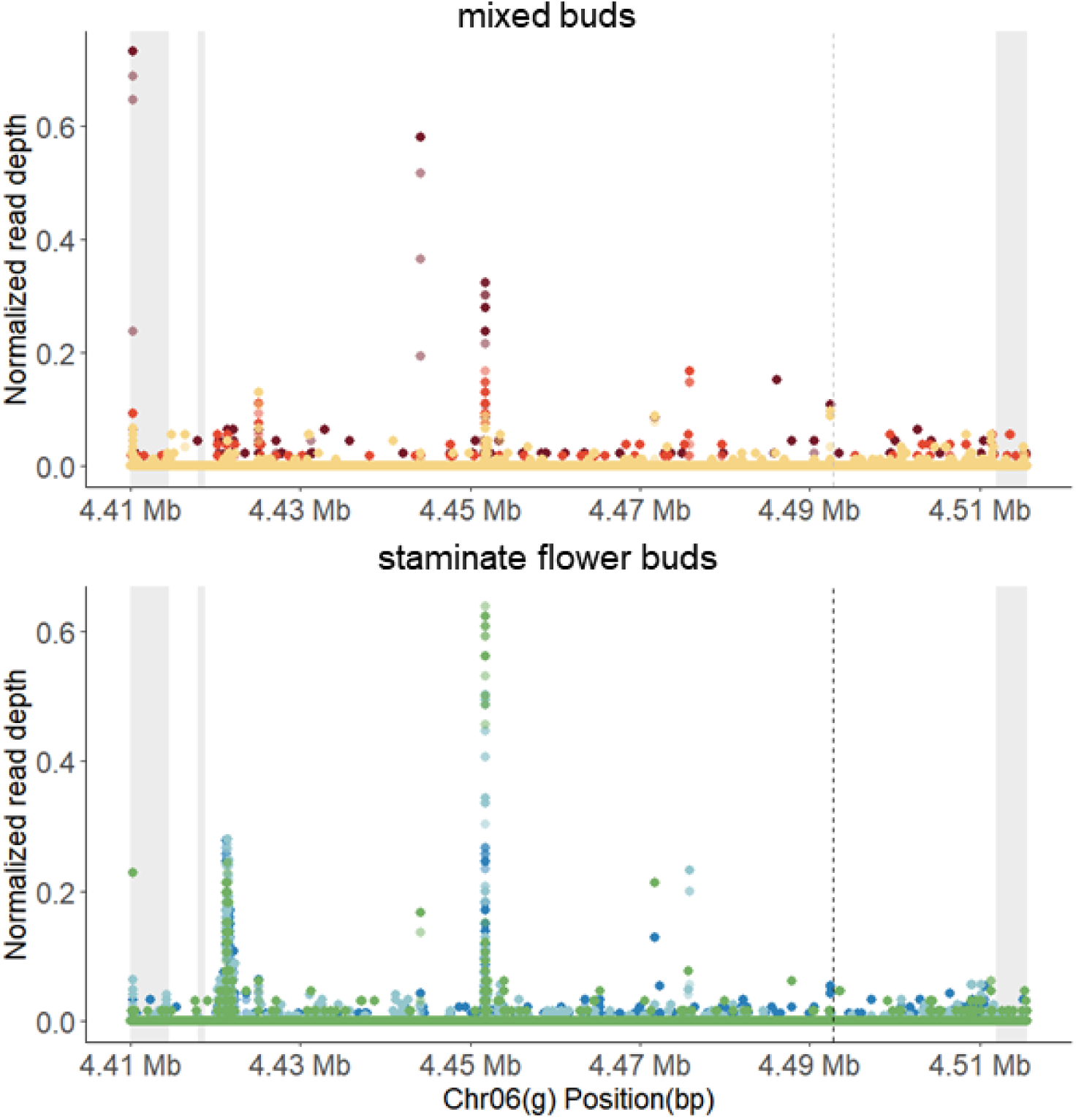
Dot plots showing Tissue-specific small RNA expression from the *g* locus. Staminate flower buds exhibit abundant sRNAs transcribed from the *G*-allele repeats, with two distinct peaks corresponding to homologous regions on the *g* allele. In contrast, mixed buds show minimal sRNA expression.

**Figure S5.**
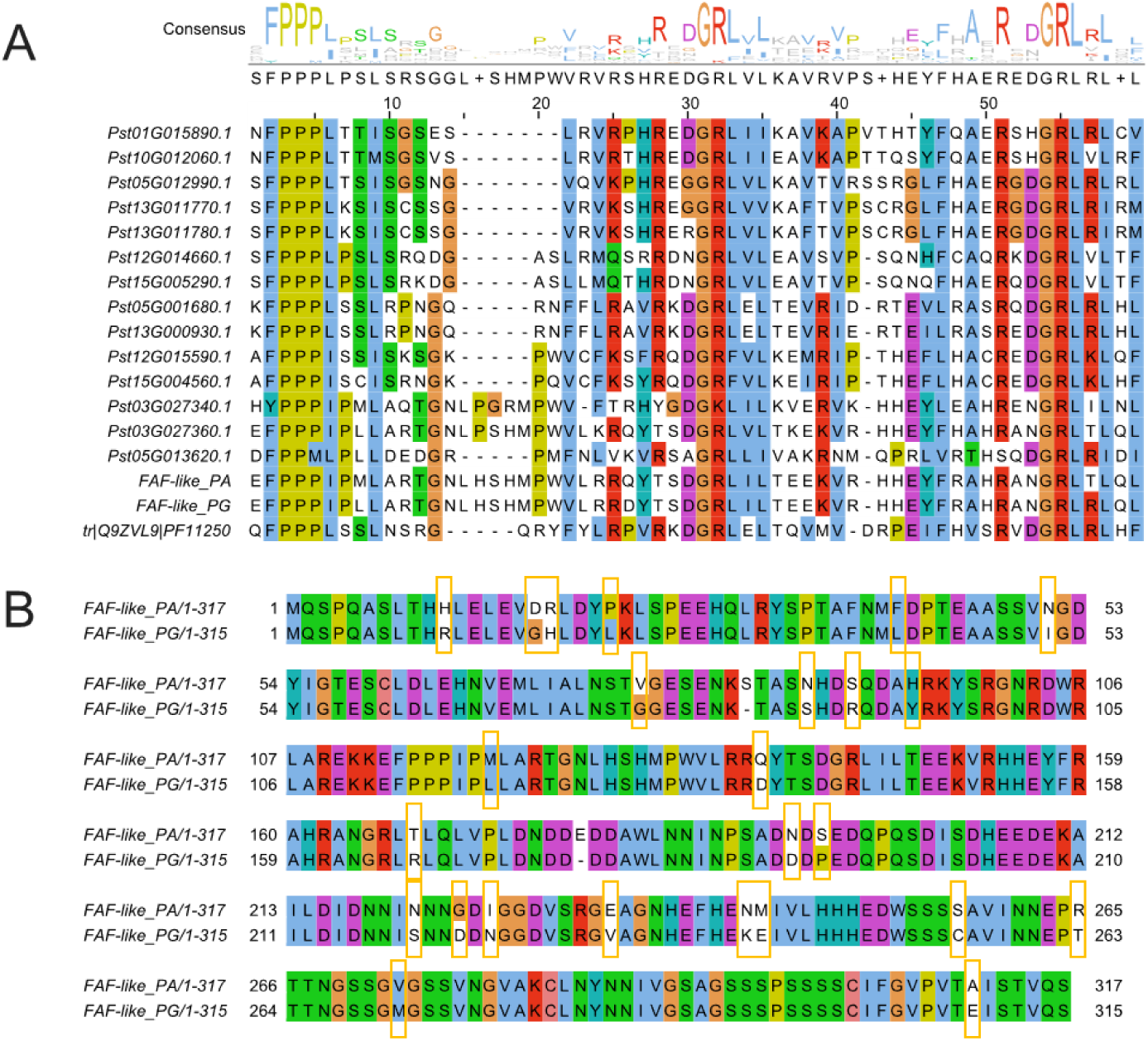
Primary structure and allelic characterization of FAF proteins in *Pterocarya stenoptera.* (A) Conserved domain architecture of *FAF* family. We identified 14 additional FAF family members in *P. stenoptera*. Multiple sequence alignment of *FAF* homologs revealed the conserved PF11250 domain (derived from *Arabidopsis thaliana* Q9ZVL9). Alignment was performed using CLUSTAL Omega with default parameters. (B) Haplotype-specific sequence divergence analysis. Comparative analysis of PA and PG haplotype-specific *FAF-like* proteins identified 25 non-synonymous substitutions.

**Figure S6.**
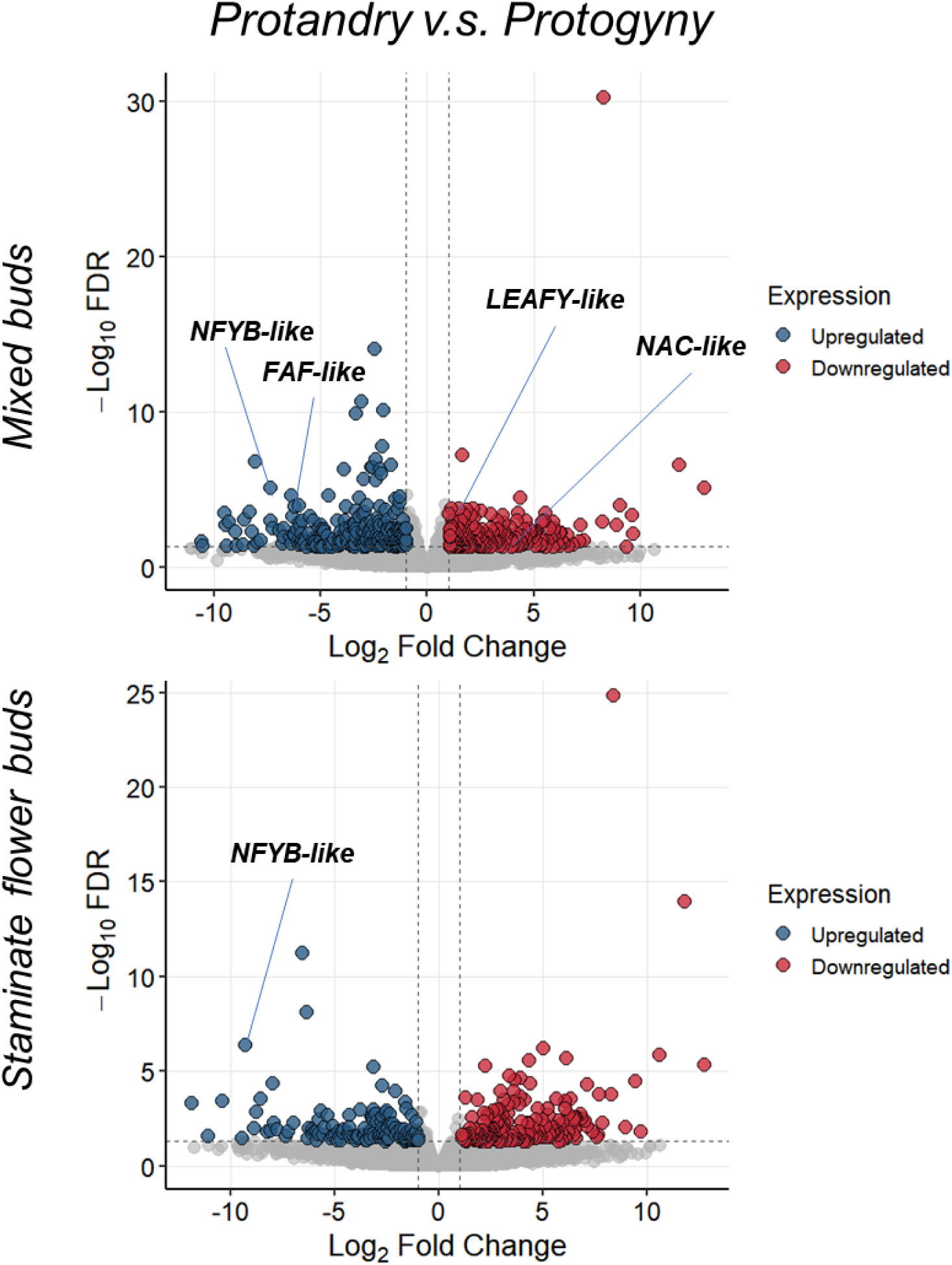
Comparative transcriptomic analysis of PA (*gg*) and PG (*Gg*) floral buds. Volcano plot of differentially expressed genes between PA/PG staminate flower buds and mixed buds (FDR < 0.05, |log2FC| > 1). Four candidate genes (*NFYB-like*, *FAF-like*, *LEAFY-like* and *NAC-like*) are highlighted.

**Figure S7.**
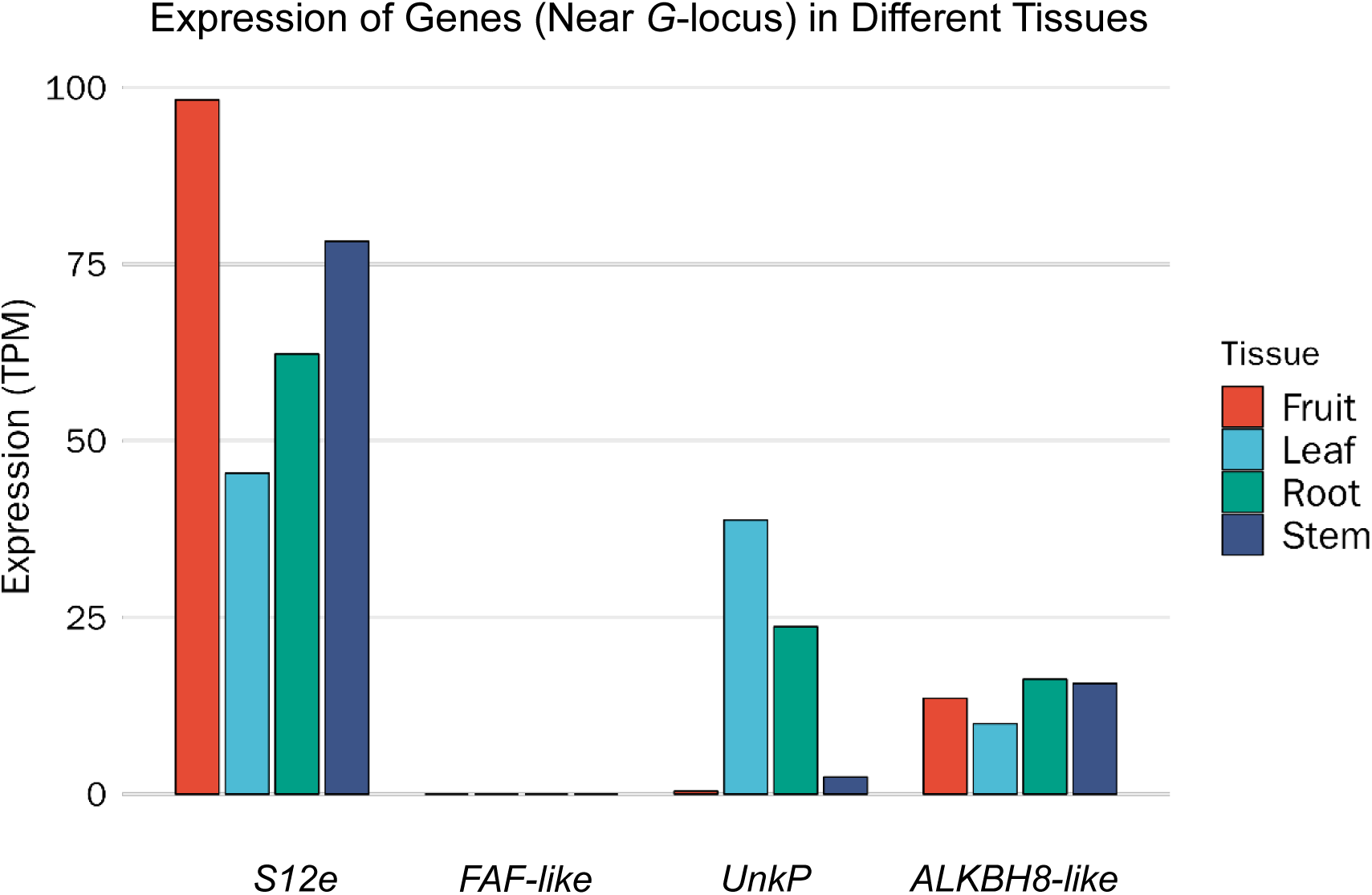
Tissue-specific expression of genes near *G* locus in *Pterocarya stenoptera*. The *FAF-like* gene at the *G* locus has no detectable transcripts in other tissues (fruit, leaves, roots or stems). Transcriptome samples from Zhang, et.al, Plant Diversity, 2025.

**Figure S8.**
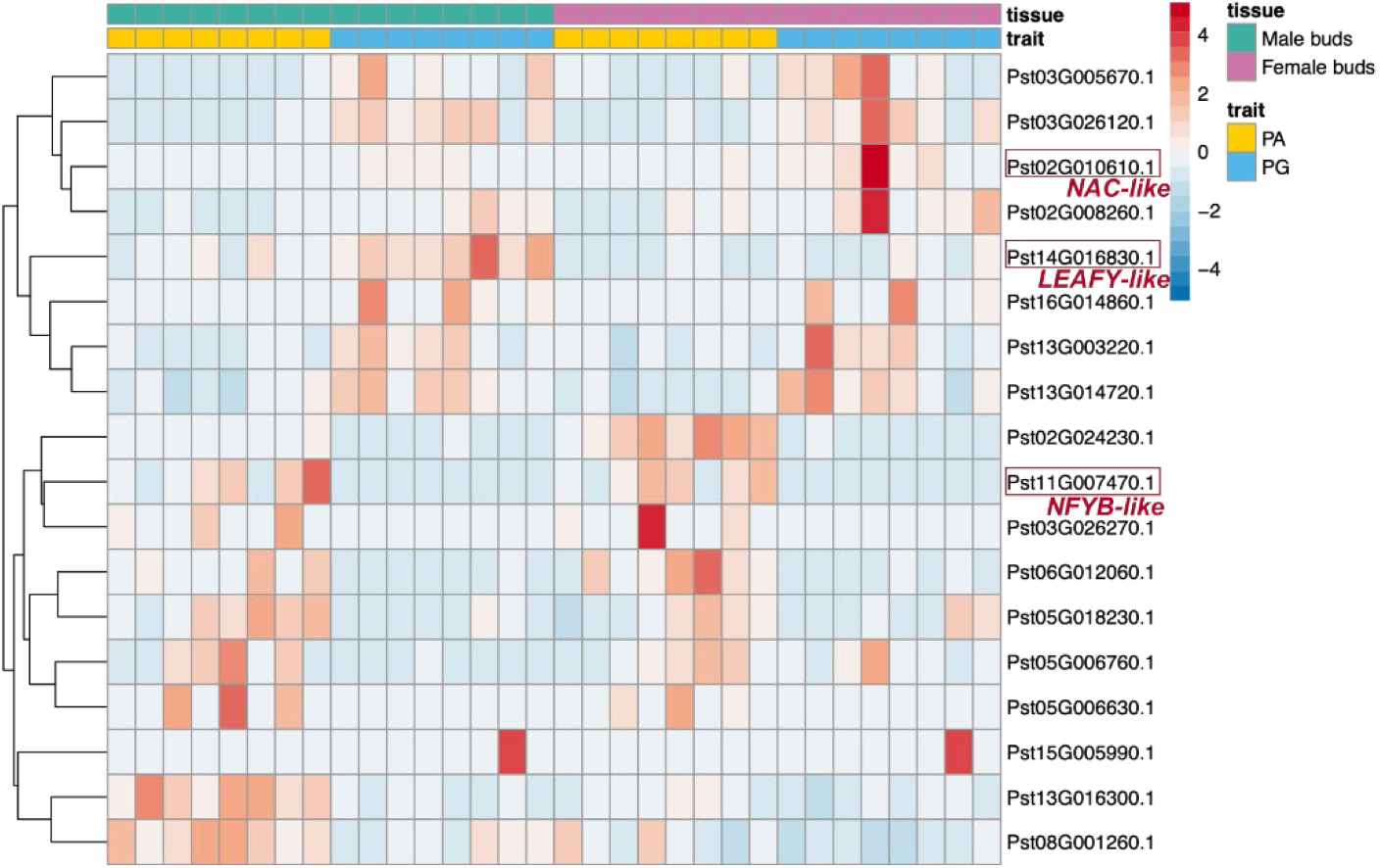
Differential expression patterns of small RNA-targeted genes in staminate flower buds. Heatmap displays 18 differentially expressed genes identified by integrated sRNA-seq and transcriptomic analysis (nine upregulated and nine downregulated in PA vs. PG, FDR < 0.05)

**Figure S9.**
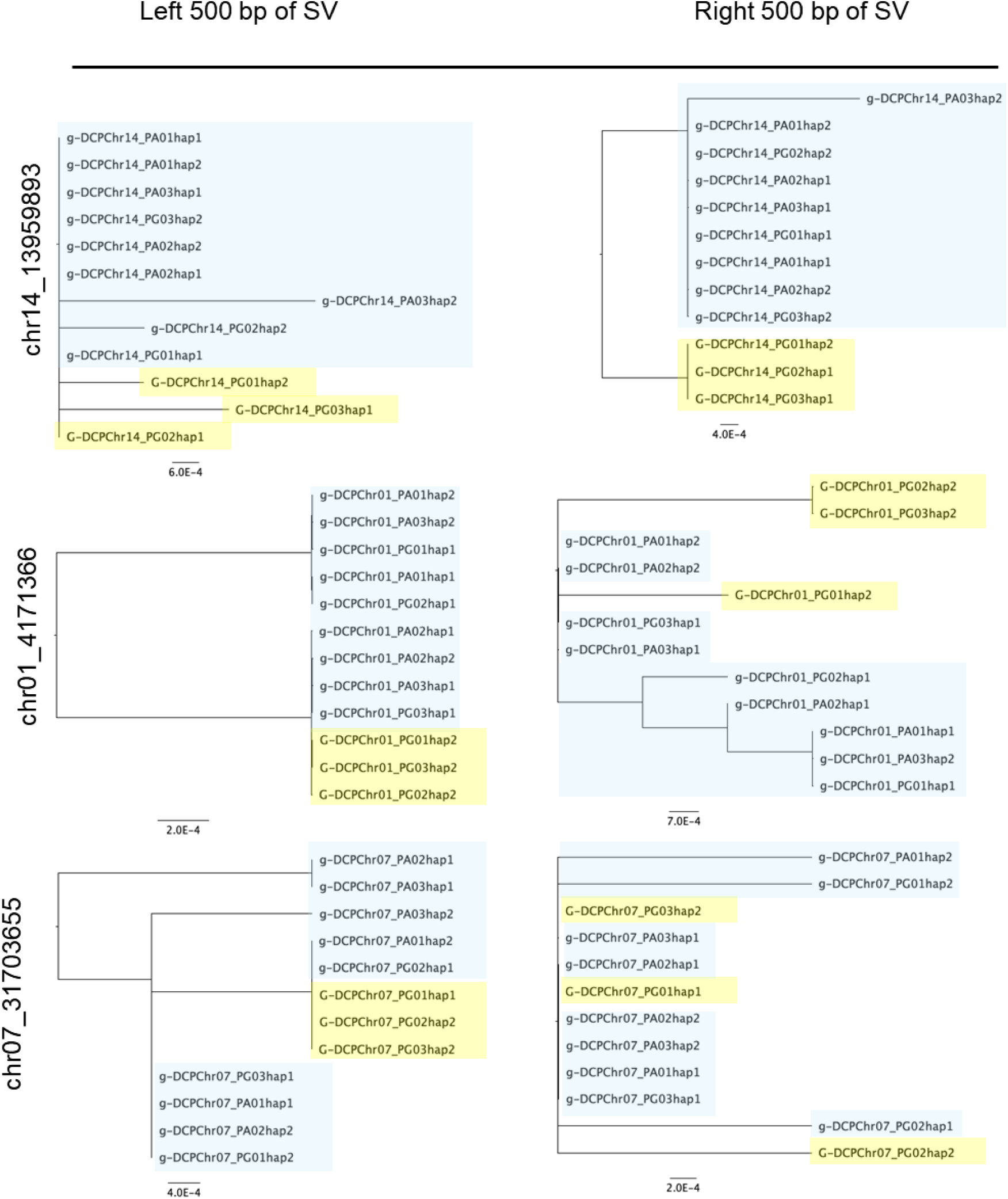
Haplotype-resolved phylogeny using 500 bp sequence flanking the candidate SVs (chr14: 13959893 bp, chr01: 4171366 bp and chr07: 31703655 bp).

**Figure S10.**
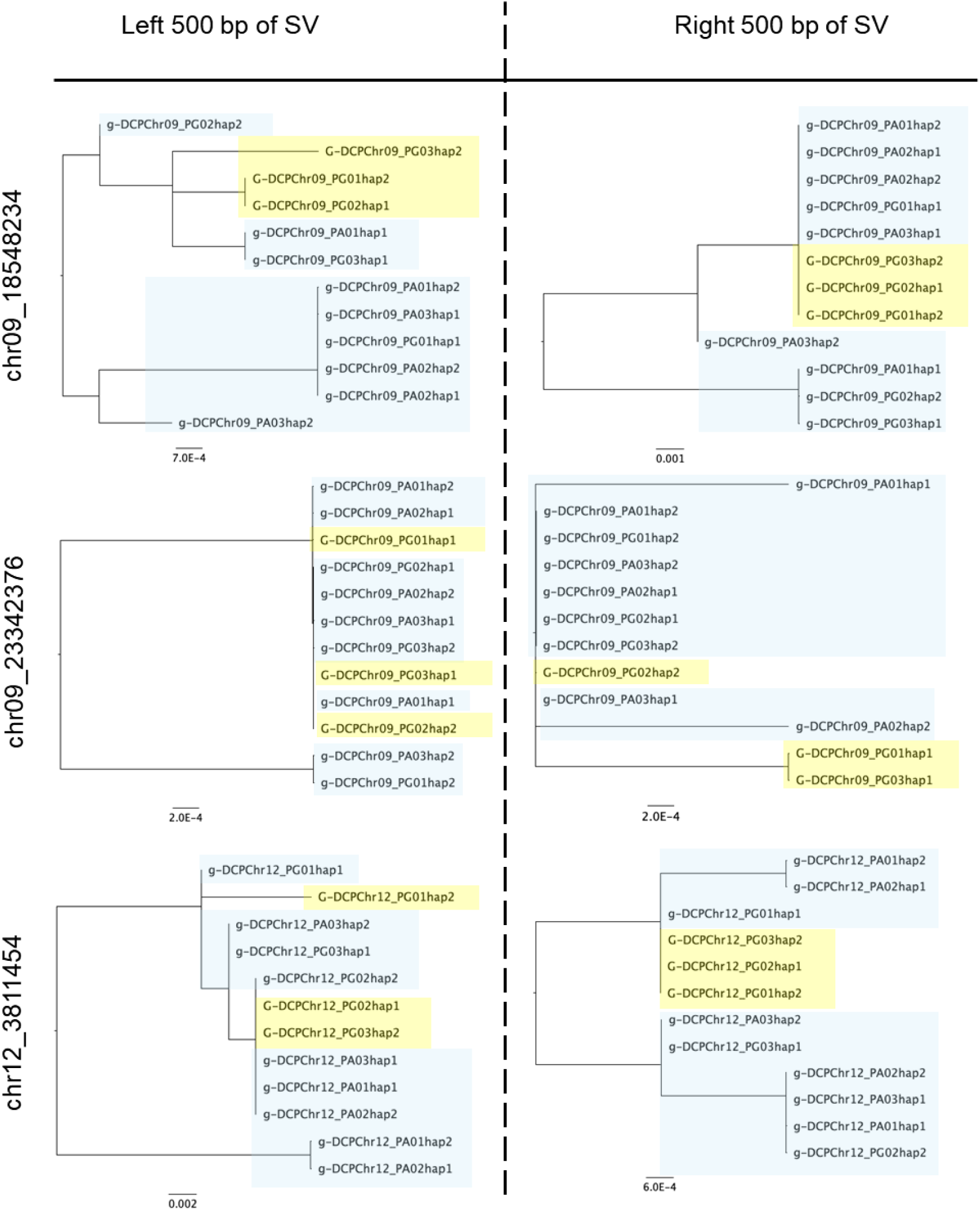
Haplotype-resolved phylogeny using 500 bp sequence flanking the candidate SVs (chr09: 18548234 bp, chr09: 23342376 bp and chr12: 3811454 bp).

